# “The Free Association Sessions”. Perspectives on a Novel Teaching Platform by Final Year Medical Students and Basic Specialist Trainees in Psychiatry

**DOI:** 10.1101/2022.11.21.517328

**Authors:** John McFarland, Gurjot Brar, Peter Hayes, De Mohamed Elhassan Abdalla

## Abstract

**Introduction:** Notwithstanding the many advantages of outcomes-based education within Psychiatry placements in Medical School and Basic Specialist Training within the College of Psychiatry of Ireland, there is limited protected time for broad Reflective Practice that appreciates the complexity of working in the Mental Health setting. Furthermore, there are limitations to the current model of restricting Reflective Practice sessions to the Balint Group Format.

**Methods:** A novel programme of structured Reflective Practice was offered to students in the School of Medicine in The University of Limerick and trainees in the Mid-West Deanery. Six student and fourteen trainee participants were subsequently invited to provide perspectives on the programme via Focus Groups. The study employed an inductive latent phenomenological approach for analysis of qualitative data.

**Results:** Five major themes emerged: These related to the teaching environment, personal identity, complexity, awareness of cognitive dissonance and the structure of the sessions. There were a number of different perceptions, relating to the participants’ stage of training.

**Conclusions:** There was evidence that the structured reflective sessions created a comfortable environment, addressed hierarchy issues and facilitated wide-ranging opportunities for reflective practice, with an observed increased appreciation for complexity in Psychiatry. There was apparent tension between controlling *content* and facilitating the *form* of group process. Nonetheless, the structure appeared more approachable for medical students and those early in training.

## Introduction

Medical students in the School of Medicine (SoM) in the University of Limerick (UL) undergo a curriculum based around twenty-six distinct Case-Based Learning (CBL) scenarios. There is a distinct focus on defined Learning Outcomes (LOs), which are consciously largely neurobiological in content.

Similarly, Basic Specialist Trainees (BSTs) in Psychiatry in Ireland progress to Higher Training after meeting a large number of LOs that are vigorously overseen by the College of Psychiatry of Ireland (CPsychI). This process is underpinned by regular supervision with Consultant supervisors and Work Place Based Assessments (WPBAs) that are reviewed centrally by the College via an annual review of progress (ARP) panel.

### Limitations of current training

Notwithstanding the many advantages of outcomes-based education, the focus on LOs has potential reductionist pitfalls:

Psychiatry is among the most complex of all medical specialities, in terms of the lack of clear categorical demarcations of conditions, the interplay with numerous psycho-social components and the interplay between the subjective views of the patient and the (highly variable) “objective” evaluation of the clinician. It is notable that even at the level of “Advanced training” for senior trainees, (Hughes *et al*. 2002) there is a predominate focus on medical aspects of illness. This contemporary push towards what Dewy refers to as “technical rationality” (Rodgers 2002) has deviated psychiatric training from the “professional artistry” (Schön 2017) that underpins the complexities of work in the Mental Health Sector.

From this background, many of the non-”core” aspects of Basic Specialist Training in Psychiatry that fall outside the strict biopsychosocial paradigm are off-loaded on to WPBAs. Nonetheless, there is evidence that many clinical supervisors have limited training for their role in these assessments (Fitch *et al*. 2008; Julyan 2009) and that the weekly timetable is not adhered to consistently (Hariman *et al*. 2020), often as the result of time-constraints.

Perhaps more concerning is that many WPBAs are designed for strict adherence to standardised procedures and are therefore limited in their scope for reflection (Simmons 2013). This deficit contrasts with supervision within the field of clinical psychology (Oyebode 2009), which may focus on areas such as ambiguity and cognitive dissonance. The Medical School approach to introducing Reflective Practice has largely been limited to the reflective writing that is incorporated into undergraduate psychiatry placements (Chaffey *et al*. 2012). It has been observed that “many” students have difficulty with reflective writing (Raw *et al*. 2005) and that those less naturally adept at reflection are more negative about the process (Rees and Sheard 2004) (Beylefeld *et al*. 2005). As such, there are potential dangers that those less open to reflection may acquire a more negative view of psychiatry if it is assessed formally.

Similarly, with respect to BSTs, The CPsychI ensures that reflective writing is incorporated into WPBAs but limited to one piece every 12 months. Again, input and critical appraisal by supervisors may be inconsistent (Julyan 2009) and these notes may often more accurately be described as “Descriptive Reflection” (Bekas 2013) as a description of events with elements of reflection.

### Balint Groups

The most substantive model of structured reflective practice in Psychiatric Training, in Ireland and the UK, is the Balint Group (BG) (Van Roy *et al*. 2015). The BG is a Reflective Practice group for clinicians focusing on psychodynamic aspects of clinical cases in order to emphasize the importance of the use of emotion and personal understanding in the doctor’s work and the therapeutic potential of the clinician-patient relationship. They are considered mandatory training for Basic Specialist trainees (BSTs) in Psychiatry in Ireland (Fitzgerald and Hunter 2003).

There is evidence that BGs promote self-reflection and gaining insight into self and patientcare dynamics as demonstrated by the Jefferson Scale of Empathy and Brief Resilience Scale (McManus *et al*. 2020). It has been reported that three BGs helped to reduce trainee doctors’ stress and anxiety levels and to enable a greater understanding of the patient experience (McKensey and Sullivan 2016) and the identity of the doctor (Ryding and Birr 2021). Nonetheless, other studies demonstrated that trainees often struggled to adapt and found the sessions anxiety provoking (Graham *et al*. 2009). Furthermore, sessions have been referred to as “repetitive” with similar areas of discussion being frequently revisited (Balint 1979; Omer and McCarthy 2010).

In studies of medical students, it has been demonstrated that many question the relevance of BGs with respect to their clinical training (Parker and Leggett 2012; Parker and Leggett 2014). There may also be issues related to limited clinical exposure, discomfort with self-disclosure and the limited time frames for group consolidation (Brazeau *et al*. 1998; Shoenberg and Suckling 2004).

With these limitations in mind, it has been previously suggested that “realistic, multifactorial” constructs that facilitate meaningful reflection are provided to students and trainees for discussion (Bekas 2013). These suggestions particularly emphasise historical aspects of psychiatry and their evolution (Bracken and Thomas 2010), personal identity, values and attitudes (Gibbs 1988; Bekas 2013; Bennett *et al*. 2017), diagnostic ambiguity, (Bekas 2013) and sociocultural components of reflective learning (Boud 2010). However, we are unaware of any studies suggesting a suitable format to address these areas.

### Aim

With consideration of the above issues, the Department of Psychiatry in UL proposed a structured programme of reflective practice sessions, “The Free Association Sessions”, to be delivered over a six-week period to both students and BSTs. These sessions aimed to address themes of complexity that promoted reflection. The content was consciously designed to avoid the potential limitations of BGs for Medical Students alluded to above and the concerns about repetitive themes expressed by BSTs.

The current study aims to analyse the perspectives of Medical Students and BSTs in Psychiatry with respect to the course and explore how the sessions may have contributed to an increased appreciation of Reflective Practice. We anticipated that there would be a resultant increased enjoyment of training, facilitating deep learning, (Webb 1997) along with an appreciation of sociocultural issues, identity, complexity and cognitive dissonance.

## Methods

### Context

Study participants were final-year medical students in the 4-year Graduate Entry Medical programme at the School of Medicine, University of Limerick during academic year 2021/2022 and Basic Specialist Trainees in Psychiatry in the UL Deanery.

Students and BSTs were invited to a, non-compulsory, six-session (each an hour long) programme of Reflective Practice. The content is outlined in Table 1. Sessions 2-6 utilised a video format, where participants watched a video-clip, prior to an in depth discussion of the arising themes.

**Table 1.** Summary of Sessions and anticipated discussion.

| Session | Anticipated Discussion |
| --- | --- |
| 1 - Balint Group with Consultant in the role of “The Doctor” | Addressing hierarchical issues<br>Introducing complexity in Psychiatry |
| 2 - Conspiracy Theorists | Addressing sociocultural tension<br>Social deviancy<br>Psychotic symptoms in the general population |
| 3 - Body Integrity Identity Disorder | What represents psychiatric illness?<br>Social deviancy |
| 4 - Cultural Approaches to Psychosis | Sociocultural models of psychosis<br>The Medical Model |
| 5 - Antipsychiatry vs “lived experience” | Diagnostic ambiguity<br>Interface with service users |
| 6 - Psychopathy | Social deviancy<br>Personality disorder / Mad vs Bad |

Analysing the perceptions of the teaching programme necessitated rich and detailed inquiry. As such, the study employed an inductive latent phenomenological approach. With such an inductive approach, the researchers anticipated emergent themes such as appreciation of complexity and cognitive dissonance and this added depth and specificity to the data.

Nonetheless, the framework allowed themes to aggregate outside these assumptions. The latent context added detail by demanding deeper analysis, enriching the interpretive layers.

### Ethical Approval and Process

University of Limerick Research Ethics & Governance Committee granted ethical approval for all Medical Student involvement in this project and the Research Ethics Committee of the UL Hospitals Group for BSTs. It was anticipated that JMF, as a member of the exam board, would hold a position of power in relation to the medical students and therefore GB (Clinical Tutor) chaired Focus Group discussions with this cohort. JMF conducted the same process with BSTs.

### Recruitment

All final-year Medical Students at University of Limerick and BSTs at the UL Deanery received emails via a gatekeeper to participate in semi-structured interviews. Six student participants and fourteen BSTs were recruited by voluntary participation.

### Data Collection

JMF and GB conceived the discussion prompts. GB chaired two Focus Group discussions with Medical Students and JMF with BSTs during the spring semester of 2022 (see Table 2). All Focus Groups were conducted and recorded via Microsoft Teams, which automatically provided transcription and data was reviewed by the participants to provide respondent validation and check accuracy.

**Table 2.** Study Participants.

| **Participants** | **Focus Group** | **Duration and comments** |
| --- | --- | --- |
| **Medical Students**<br>3 Female, 3 Male<br>Age range 26-30 | **Focus Group 1:**<br>3 Participants<br>Students A, B, C | 45 minute duration<br>All students contributed to discussion |
|  | **Focus Group 2:**<br>3 Participants<br>Students D, E, F | 45 minute duration<br>All students contributed to discussion |
| **Basic Specialist Trainees**<br>8 Female, 6 Male<br>Age range 28-46 | **Focus Group 3:**<br>BSTs A-G | 90 minute duration<br>All BSTs contributed to discussion |
|  | **Focus Group 4:**<br>BSTs H-N | 90 minute duration<br>All BSTs contributed to discussion |

### Data Analysis

Data was analysed using an adapted six-stage guide (Braun and Clarke 2006). The research team (JMF, PH, MEH) initially familiarised themselves with the data by reading and rereading the transcribed data. Preliminary codes where generated relating to the transcribed data and agreed upon, after discussion amongst the research team before collation into potential themes. Data sorting was performed manually with colour coding in Microsoft Word. Themes were subsequently reviewed in relation to the entire dataset and revised as required. All themes and subthemes were further categorised before the final report was produced. Thematic mind maps gave an overview of the process to the research team. Memos were kept throughout the duration of analysis. Assumptions of the researchers were identified and made explicit at the outset and subsequently revisited at points of divergence or convergence. To avoid transferring assumptions, researchers questioned and challenged each other to support reflexivity.

## Results

Five major themes emerged: environment, personal identity, complexity, awareness of cognitive dissonance and structure of sessions (see Table 3)

**Table 3.** Themes and Subthemes generated from Focus groups.

| Theme | Subthemes |
| --- | --- |
| Environment | Comfort<br>Hierarchy issues<br>Peer-Based Discussion |
| Personal Identity | The Human Doctor<br>Public Perceptions of Psychiatry |
| Complexity in Psychiatry | Challenging Reductionism<br>Thinking Outside the Box |
| Awareness of Cognitive Dissonance | Reflective Practice<br>Changed my View<br>Clinical Approaches |
| Structure of Sessions | Format<br>Agency |

### 1 Environment

#### (i) Comfort

Both students and BSTs commented on the informal nature of the sessions in positive terms:

*It feels like it’s time off, but at the same time you are learning (BST A Year 1)*

*I guess it was quite casual, people can kind of blurt out whatever they want (Student A)*

#### (ii) Hierarchy issues

Although there was some evidence that Medical Students initially found the sessions intimidating, the approach of senior psychiatrists reflecting on material appeared to have a beneficial effect on perceived hierarchical issues:

*I think there is some good in like going to a session and feeling uncomfortable in it and working yourself through that (Student B)*

*It is refreshing just to be able to have a conversation with the Doctor who has so much experience to say to you “What is your take on that? “ (Student C)*

BSTs also reported on the benefits of input at differing levels of seniority:

*It’s quite a powerful learning experience I think… to know there’s people who’ve gone through all this stuff before and…they’ve had the same kind of difficulties. It’s a very useful tool and probably something that’s underutilized (BST B Year 2)*

#### (iii) Peer-based discussion

Medical Students expressed benefits from the open discussion with peers around the sessions, highlighted deep learning benefits and appeared stimulated to continue the discussions outside the situated learning environment:

*It’s like an open discussion and you actually get to think (Student D)*

*It was a really interesting topic…you actually got some good insight as well when you talked about it outside of the session. (Student B)*

BSTs demonstrated similar benefits in terms of open discussion but also particularly appreciated support from colleagues, having opinions challenged and a therapeutic benefit to the sessions:

*Getting different perspectives on a situation suddenly can like change your whole approach (BST C Foundation Year)*

*I enjoyed the therapeutic element of it as well. The fact that we can talk and vent (BST D Year 2)*

### 2 Personal Identity

#### (i) The Human Doctor

Both students and BSTs commented on potential benefits from the sessions in terms of self-reflection. With increased clinical experience, BSTs demonstrated the capacity to relate this to both self-care and interactions with patients:

*The sessions were very good at, like, becoming introspective…looking at yourself (Student B)*

*(Discussing) feelings that would be attached to a patient encounter… that’s the main thing that I’ve taken away from the sessions is that it’s actually very beneficial for yourself and for the care of patients that you actually reflect (BST E Year 3)*

#### (ii) Public’s perceptions of Psychiatry

Similarly, where students appreciated challenging views of psychiatry from outside the profession, BSTs related this to their clinical practice.

*Others can see the medical world defining what they do, defining normal, whatever. So I think it was interesting to see that like, that’s not people don’t always agree with that. (Student D)*

*I think in the public’s eyes, it’s very blurry and very misunderstood…It becomes very important to clarify to people that we ‘re in a science of speculations and I think when you approach it that way, that reassures the patient that we’re working on this together. (BST D Year 2)*

### 3 Complexity in Psychiatry

Both students and BSTs commented that the sessions helped them reflect on themes of complexity in clinical psychiatry:

#### (i) Challenging Reductionism

The two cohorts expressed that the sessions helped them reflect on the limitations of diagnosis:

*It’s made it more complicated in a way that like, there’s just so much more to it than you could take in a 10 or 20 minute or even hour history. (Student A)*

*The confidence to allow yourself a few sessions to get to know a person and to hold back on a diagnosis and just to let that kind of play out a little bit more and that that doesn’t mean that you don’t know what you’re doing. (BST C Foundation Year)*

#### (ii) Thinking outside the box

There was an appreciation of the need to remaining open to the multiple complexities at play outside strict psychiatric constructs of diagnosis and treatment:

*That’s an important part to kind of really understand the patient and everything that’s going on with their illness but also their lifestyle and living situation… it’s complicated… it’s not just black and white… So just to keep your mind open. (Student A)*

*Psychiatry is a field in which there’s an overlap between a very sort of strict science as well as an aspect of humanities as well, philosophy and other sort of less black and white (constructs). (The sessions) definitely increased my tolerance of ambiguity and (the) abstract nature of psychiatry. (BST F Foundation Year)*

### 4 Awareness of Cognitive Dissonance

#### (i) Reflective practice

There was evidence that the sessions increased students’ awareness of the benefits of Reflective Practice and similarly for BSTs in terms of challenging their thoughts on an ongoing basis:

*In medicine at the moment… to challenge how you think is so important, especially before we start into a job where that’s going to happen every day. (Student C)*

*So in psychiatry, it’s been a big adjustment to kind of sit with that uncertainty. (BST G Year 1)*

#### (ii) Changed my view

Both groups appreciated that the course material had challenged their views and had influenced their perspectives:

*We need to be comfortable sitting in that position…just because someone doesn’t like your profession doesn’t mean that you can’t treat them or, like, do the best you can for them…kind of tiptoeing around someone. (Student B)*

*I think there is a case that comes and just changes…that we go through an epiphany moment of changing our whole perspective in a certain area in the field. (BST D Year 2)*

#### (iii) Clinical approaches

There was evidence that the sessions became an aid for Medical Students on clinical placements and that BSTs found the reflective groups had an impact on their clinical practice: *I was not prepared for the wild answers. So I think the freeform sessions probably helps with that because…you’re listening to someone on a video and they’re saying very wild things, whereas like in the (formal teaching) they are very more classic cases. (Student E)*

*I might go into the clinic the next day and I might approach things slightly differently or… I might explore something that I probably hadn’t explored before so (the teaching programme) was quite stimulating. (BST B Year 2)*

### 5 Structure of Sessions

#### (i) Format

Medical Students appreciated the link between the Reflective Practice sessions and theoretical teaching:

*The fact that it led from what we did earlier on in, in the lectures and just helps an awful lot. To not only solidified the information, but look at how you would apply that. (Student D)*

Interestingly, BSTs at an early stage of training found the structured approach to Reflective Practice useful, whereas those approaching HST expressed a preference for formal Balint groups:

*(Using) a specific presentation and having a critical discussion about this it’s way more interesting, but it’s also it’s much better learning. (BST G Year 1)*

*(When asked were structured sessions more useful than Balint Groups) Not really -I suppose common things are common, so they will almost definitely be useful. (BST H Year 4)*

#### (ii) Agency

Notably, the sessions appeared to result in an increased sense of agency from both cohorts, who expressed a wish to participate in the evolution of the teaching sessions:

*It would be useful for the freeform sessions to lead on from what we discussed (had been taught) before. (Student F)*

*Have a list of potential topics or ask your audience before starting the next session… rotation on what topics they would like to have. (BST I Foundation Year)*

## Discussion

The perspectives on the Free Association sessions, from medical students and trainees, fell into five main themes. These related to the environment, personal identity, complexity, awareness of cognitive dissonance and structure of sessions.

There were differences in how students perceived the sessions compared to trainees and within trainee cohort, some differences relating to their seniority.

### Environment

A universal view was the benefit of a comfortable, casual environment to discuss topics.

There was evidence that this setting facilitated deep learning (Webb 1997) within the medical student cohort and eased concerns centred on hierchial obstacles:

Reflective Practice groups have been observed to evolve over time and this is consistent with a Community of Practice (CoP) model (Austin and Duncan-Hewitt 2005) where learning involves a process of socialisation in which newcomers to a group move from peripheral to full participation, eventually becoming part of the shared practices and beliefs common to that group process. The discomfort that medical students have previously expressed in the BG literature (Parker and Leggett 2012) is part of this process. Although students in the current study expressed initial anxiety, it was apparent that the structured themes of the sessions aided medical student participation and there was appreciation of the consciously flattened sense of hierarchy in the format.

Within the BST cohort, there was evidence that the slow process of what Wenger termed ‘legitimate peripheral participation’, (Wenger 2011) was expedited. Nonetheless, as discussed *vide infra*, our data suggests a different appreciation of reflection as training progressed amongst trainees.

Acknowledgement of this process is central: Medicine historically has a significant hierarchical structure and students and trainees often envisage challenging sociocultural associations with seniors. There is a complexity with this impression of hierarchy within Reflective Practice groups. It may be perceived that “higher-order” reflection is appropriately restricted to those with increased seniority and, unless this is addressed, it may discourage an innate capacity for reflective thinking in students or trainees. An interesting analogy comes from industry where it was observed that lack of explicit attention to power relations within CoP theory may lead to surreptitious control of groups (Gee 1996).

As such, the approach at outset of conducting a Balint session, with a consultant taking the “doctor” role, appeared to be effective. The perspectives were consistent with previous constructs that used the observation of seniors “struggling” with complexity as a tool to “flatten” the educational environment (Dempsey and Kauffman 2017).

To further counteract hierarchical issues, our sessions explicitly emphasised the collective and shared nature of learning (Lave 2009). There was evidence that the sessions merged learning and practice, allowing for a meaningful reflection in the context of participation and assimilated the construct of *reification* within a team based discussion.

Our data demonstrated that this iterative process led to metacognitive activities, where evolving attitudes and positions reached a dialectic relationship with practice; “reflection-on-action” leading to “reflection-in-action”(Schön 2017). A central tenant of the relationship between Reflective Practice Groups and CoPs is the circular process of being shaped by experiences and in turn addressing how the group shapes the (educational) environment. As such, a further intended outcome would include an increased sense of agency and position within the aforementioned hierarchy. In this respect, it was pleasing to see that as training progressed, participants were more likely to suggest topics and developed an increased sense of initiative. Of note, this was also evident within the medical student cohort. Although not directly assessed in this study, this process was further facilitated by a focus on “vertical integration” (Austin and Duncan-Hewitt 2005) where trainees became involved in student sessions.

### Identity

The observation that our Reflective Practice sessions helped foster an awareness identity is consistent with previous work with BGs (Ryding and Birr 2021). Notably, a survey of tutors in psychiatry found that the typical desirable abilities of a trainee psychiatrist should include overall clinical competency in diagnosis, investigations and management (Bhugra *et al*. 2009). It is of interest that this large study had a primary focus on what the psychiatrist *does*, rather than the psychiatrists’ *identity*. Although areas of professionalism are addressed in the psychiatric curriculum, at both medical student and trainee level, there is a lack of focus on the value system of psychiatrists. It is logical that areas such as empathy and understanding may have considerable impact on the psychological distress of a patient. This places the “character of the practitioner” (Radden and Sadler 2010) in a position of central importance with respect to the ethical practice of psychiatry. Indeed, the importance of the “person” of the psychiatrist, has historically been given equal credence to their role as a technician(Menninger 1952).

Although questionable, this assertion is consistent with the theme of identity that was apparent within the groups. The shift in the content of the sessions towards the humanities and reflection on the “art” of psychiatry appeared to be an enjoyable element for both cohorts, with a potential role in both attracting a diverse population of medical students to psychiatry and producing well-rounded practitioners (Brakoulias 2014).

Furthermore, in terms of Medical Students, previous observations that “the ability to think reflectively… developed only after some practice experience” (Mann *et al*. 2009) appeared to be counteracted by the structured sessions and strengthened the argument that early reflection on identity may work towards self-awareness in this group (Bennett *et al*. 2017).

### Complexity

The theme of “thinking outside the box” stood out in both student and BST observations. Commentary was congruent with the basis that clinical psychiatry is commonly less precise than other branches of medicine. Trainees and Medical Students demonstrated an appreciation of complexity and the need to be considered in their decision-making, acknowledging the observation that clinical practice is both “complex and high stakes” (Hawgood *et al*. 2008).

Although the Colleges of Psychiatry in Ireland and the UK have responded to any overtly “medical” approach by advocating the “biopsychosocial model”(Engel 1981; Ghaemi 2009), it has been increasingly acknowledged that psychiatry requires “complex professional decisions that draw on diverse, and sometimes competing, fields of knowledge” (McFarland *et al*. 2009; Baker 2012) and employs a number of disparate intellectual approaches (Bhugra 2010; Smyth *et al*. 2021). As such, the observations in the current study are consistent with opinion that the biopsychosocial model in itself is overly simplistic (Roberts and Wolfson 2004) and that further dimensions, including spiritual, philosophical and cultural aspects, should be considered in the holistic care of patients with mental illnesses (Bhugra *et al*. 2009). Within this wide range of complexities and considering under-resourcing of many mental health teams, psychiatry has been referred to as “an impossible profession” (Bloch 1997).

With this background, there may be a lack of common ground between outcomes-focused educational content for psychiatric training and *de facto* clinical practice; the “widening gap” that Schon observed between theory and practice (Schön 2017). Our sessions provide a possible template for narrowing this gap and exploring the complexity of “coal-face” clinical psychiatry in a structured teaching programme.

### Cognitive Dissonance

The “thinking outside the box” theme, logically extended into areas of cognitive dissonance, where previously unexplored approaches influenced trainee’s clinical practice and challenged students’ perceptions of treatment. It was apparent that there was an appreciation of conflicting cultural and philosophical bases for the assessment of patients, which has previously been observed as an under-utilised area within psychiatric training (Aftab and Waterman 2021). Indeed, a reflective approach to differing sociocultural views of the complexities of illness definition is an evolving necessity for the modern psychiatrist (Roberts and Wolfson 2004). Our programme suggests that structured reflection would aid the development of basic philosophical tools to address these challenging areas from a position of respect and appreciation of complexity (Horien and Bommersbach 2021).

### Structure

Previous work has raised concerns that high-fidelity BGs may become repetitive with similar themes arising (Omer and McCarthy 2010). Furthermore, for Medical Students, there may be limited clinical exposure to cases suitable for discussion. With this in mind, it is unsurprising that students were positive with respect to the structured format of the sessions. Interestingly, as training progressed, views became more positive towards high-fidelity Balint, in contrast to those in Foundation Year or earlier stages of BST training who were appreciative of a structured format. It has been suggested that attempts to “manage” CoP’s are paradoxical as the perceived benefits of the model relate to spontaneity and informality (Austin and Duncan-Hewitt 2005). This also relates to BGs, where the facilitator has an essentially passive role (Omer and McCarthy 2010) and this apparent tension between controlling *content* and facilitating the *form* of group process is an important avenue for consideration.

It does seem apparent from the observations in the current study that, if they are to be of value to junior trainees and students, reflective practice groups require anchoring to both a structure and an evidence base (Barwick *et al*. 2009). Nonetheless, to retain the philosophy of the sessions, control over the content must allow for genuine reflection.

## Conclusion

In addressing the seemingly paradoxical constructs of content control and meaningful reflective practice, our programme provides some preliminary evidence that structured reflective sessions created a comfortable environment, addressed hierarchy and facilitated wide-ranging reflection on multiple areas. Furthermore, the structure appeared more approachable for medical students and those early in training. Future studies may utilise similar templates to explore how Reflective Practice is introduced to those with limited exposure to clinical psychiatry.

### Limitations

**Table 4.**
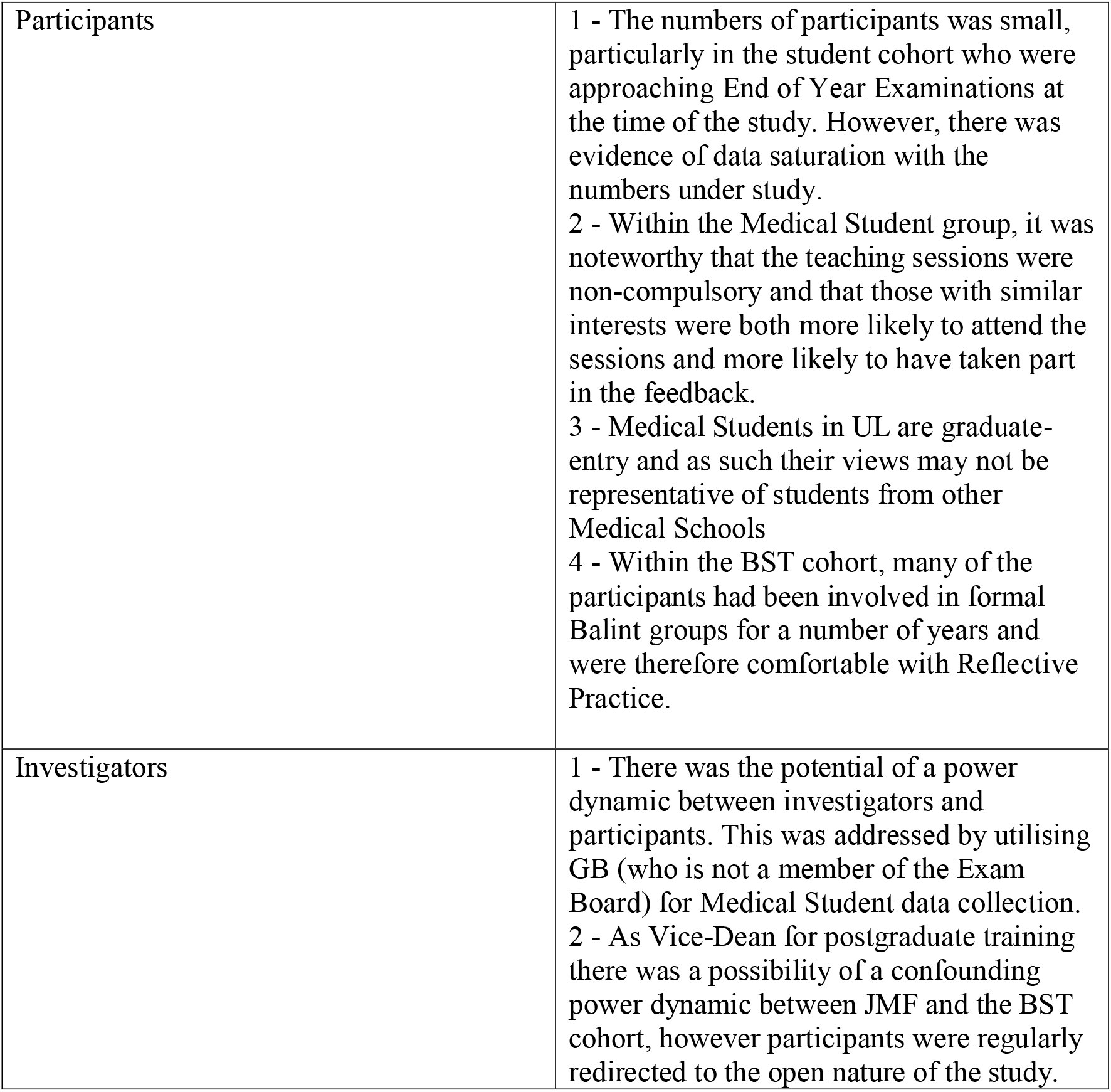
Limitations of Study.

|  |  |
| --- | --- |
| Participants | 1 - The numbers of participants was small, particularly in the student cohort who were approaching End of Year Examinations at the time of the study. However, there was evidence of data saturation with the numbers under study.<br>2 - Within the Medical Student group, it was noteworthy that the teaching sessions were non-compulsory and that those with similar interests were both more likely to attend the sessions and more likely to have taken part in the feedback.<br>3 - Medical Students in UL are graduate-entry and as such their views may not be representative of students from other Medical Schools<br>4 - Within the BST cohort, many of the participants had been involved in formal Balint groups for a number of years and were therefore comfortable with Reflective Practice. |
| Investigators | 1 - There was the potential of a power dynamic between investigators and participants. This was addressed by utilising GB (who is not a member of the Exam Board) for Medical Student data collection.<br>2 - As Vice-Dean for postgraduate training there was a possibility of a confounding power dynamic between JMF and the BST cohort, however participants were regularly redirected to the open nature of the study. |

